# Targeted neutralizing IgY antibodies against SGLT1 glucose transporter reduce glucose uptake and improve glycemic profile in vivo

**DOI:** 10.1101/2023.05.06.539723

**Authors:** Uday Saxena, Kranti Meher, RN Arpitha Reddy, K Saranya, Gopi Kadiyala, Subramanian Iyer, Subrahmanyam Vangala, Satish Chandran

## Abstract

Despite the use of several drugs available to treat type 2 diabetes, many patients are unable to reach their target fasting plasma glucose and HbA1c levels. SGLT1 is the major intestinal transport transmembrane protein which functions in uptake of dietary glucose. If we antagonise the binding of dietary glucose to this transport protein, it is expected that blood glucose lowering will follow. We designed specific inhibitory avian antibodies (IgY) against the extracellular glucose binding domain of SGLT1 and tested their potential in glucose lowering. We demonstrate here the antibodies block uptake of glucose and improve the glycemic profile in vivo and represent a novel approach to inhibiting dietary glucose absorption as treatment for diabetes.

## Introduction

Pre-diabetes and Type 2 diabetes prevalence is increasing worldwide. Pre-diabetes is the period of above normal blood glucose levels prior to the onset of full-fledged Type 2 diabetes. It lasts for a few years before transitioning to full blow diabetes and is potentially reversible. Gestational diabetes is often seen during second trimester of pregnancy and has to be managed very carefully since drugs are not an option in this situation. While there are many drugs for Type 2 diabetes, patient compliance can be low, and medication is often is not sufficient to meet target glucose levels.

Conventional medications prescribed for diabetes include insulin, insulin analogs, sulfonylureas, metformin, DPP4 inhibitors and GLP-1 analogs. Some of these drug interventions can be accompanied by adverse side effects such as hypoglycemia, lactic acidosis, gastrointestinal, weight loss as side effects. Therefore, a need exists for new acceptable, safe, and effective treatments for the lowering of blood glucose levels as an anti-diabetic treatment for pre-diabetic diabetic patients and gestational diabetes.

Sodium–glucose co-transporters SGLT, members of the solute carrier family SLC5, are high-affinity Na+/glucose co-transporters. SGLT1 transports glucose and galactose across the luminal (gut) side of enterocytes and is the first step in the absorption of sugars such as glucose and galactose from diet (1). In the kidney, SGLT1 is located on the apical (urine) side of the proximal tubule and facilitates the reabsorption of urinary glucose from the glomerular filtrate along with SGLT2 which is predominant family member in the kidney. SGLT1/SGLT2 have a key role in the absorption of glucose in the kidney and/or GI tract.

Upto 70% of the blood glucose could be derived thru the diet, therefore inhibition of intestinal SGLT1 function is an exciting new approach to glucose lowering anti-diabetic concept. SGLT1 is strongly expressed in the apical brush border of the small intestine and the late proximal tubule of the kidney, where it is critical for absorption/reabsorption of glucose into the blood stream (Figure 1). SGLT1 inhibition will lower blood glucose in diabetic patients by reducing dietary glucose absorption in the intestine as well as by increasing the release of gastrointestinal incretins like glucagon-like peptide-1. In short-term studies, inhibition of SGLT1 and combined SGLT1/SGLT2 inhibition appeared to be safe and not associated with increased rates of hypoglycemia or clinically relevant gastrointestinal side effects (2,3,4).

**Figure 1.**
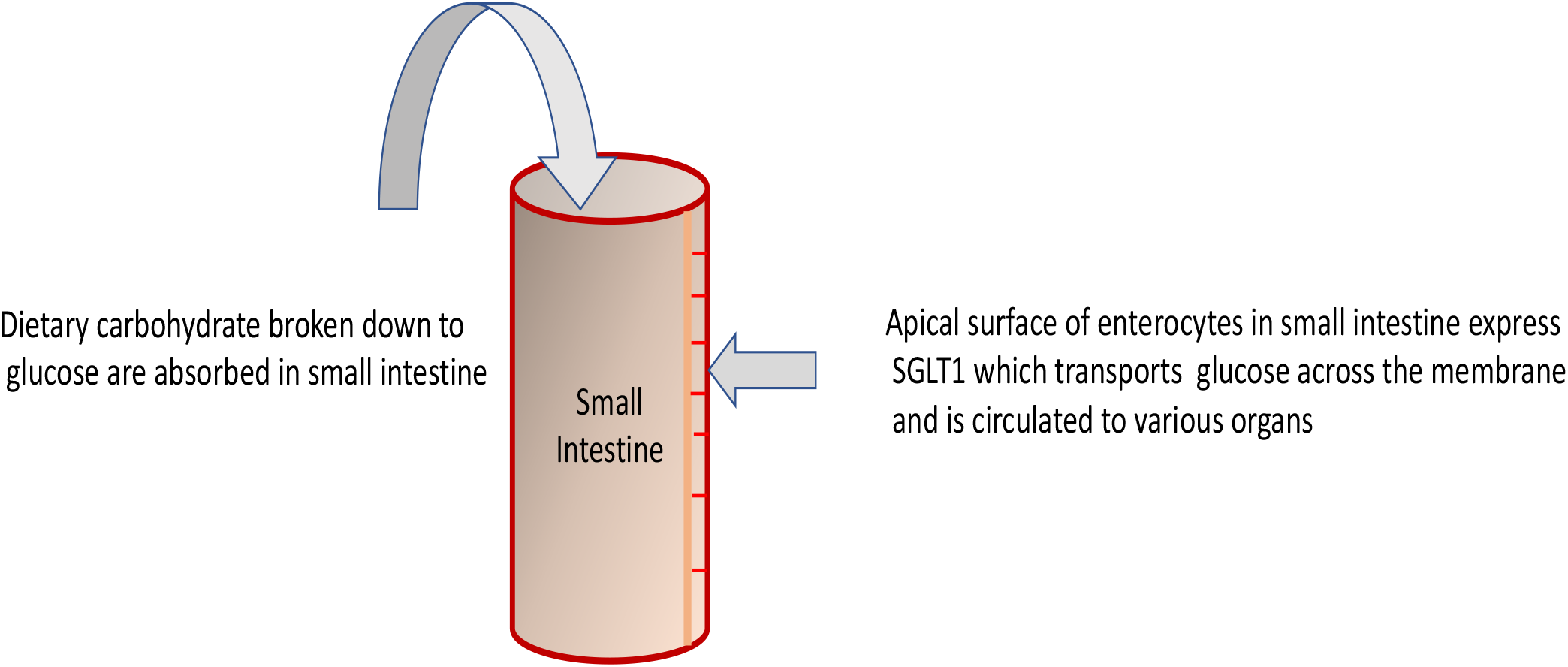

As mentioned above, SGLT1 is mainly expressed on the apical membrane (i.e., the gut side) of enterocytes and absorb dietary glucose (Figure 1). SGLT1 is a 75-kDa membrane protein with 14 transmembrane α-helices, an extracellular amino terminus and an intracellular carboxyl terminus (5,6,7). We hypothesized that neutralising antibodies against the extracellular glucose binding portion of SGLT1 may reduce glucose uptake and improve glycemic profile after oral glucose load. Here we show antibodies directed against the extracellular section of this protein inhibits glucose uptake in human intestinal immortalized cell line Caco2 and improves glycemic profile in vivo.

## Methods

### Caco2 assay

Glucose uptake studies using Caco2 cells were performed using a typical glucose uptake assay. Briefly Caco2 cells, (NCCS Pune, India) were maintained in Dulbecco’s modified Eagle’s medium (DMEM) supplemented with 10% Fetal Bovine Serum (FBS) and 100 μ g/mL penicillin and streptomycin, at 37 °C in an atmosphere of 5% CO2.2-NBDG glucose uptake. 2-(N-[7-nitrobenz-2-oxa-1, 3-diazol-4-yl] amino)-2-deoxyglucose (2-NBDG) (Molecular Probes-Invitrogen, CA, USA) was used to assess glucose uptake in Caco2 cells. Cells were kept in glucose free medium for half an hour before insulin stimulation. Cells were pre-treated the SGLT1 peptide or the anti-IgY antibodies first. Cells were stimulated with 100 nM insulin for 10 and 20 min and incubated with 10 μ M of 2-NBDG for 15 min. The reaction was stopped by washing with cold PBS three times and the cells were lysed 0.1% Triton X. The lysate was then used to read florescence at 535 nM. The data is reported as percent of control cells (no treatment, incubation with media alone).

### Preparation of anti-SGLT1 IgY

Anti-SGLT1 IgY used here were prepared using a protocol similar to that described before (9) except that the SGLT1 peptide used as antigen was used at concentrations > 10 mg/animal.

### ELISA protocol

The ELISA protocol used here to assess the binding of anti-SGLT1 antibodies to the peptide antigen was essentially the same as described before (9).

### Oral Glucose Tolerance Test (OGTT)

Oral Glucose tolerance test in male wistaria rats was performed using the protocol similar to what we have reported previously (10,11). The studies were performed by Liveon Biolabs PVT Ltd, India, under IAEC approval. The details of this study are described below.

### Animal Acclimatization

The animals were acclimatized to experimental room conditions for a period of 6 days prior to initiation of treatment. Body weights were recorded at the initiation and end of acclimatization period. Animals were observed for mortality and morbidity once daily during acclimatization period. Veterinary examination was performed before selecting the animals and only healthy and active animals were used in the study.

### Animal Identification

During acclimatization period (Temporary identification), each animal was identified by animal number written on tail using marker pen. The cages were identified with cage cards indicating study number, study code, species, strain, sex, acclimatization start and acclimatization end date.

During treatment period (Permanent identification), each animal was identified by tail marking with animal accession number written on tail using permanent marker pen. The cages were identified with cage cards indicating study number, animal accession number, study code, species, strain, sex, treatment start date and treatment end date.

### Randomization and Grouping

Grouping of animals was performed on the fifth day of acclimatization by body weight randomization and stratification method and allocated into six groups G1 to G6. G1 and G4 control groups, G2 were the reference item group. G3, G5 and G6 were the test item groups. The body weight variation within the groups of animals did not exceed ±20% of the mean body weight.

### Study Design and Group Allocation

#### Single dose OGTT study

The animals were assigned into 3 different treatment groups as shown below

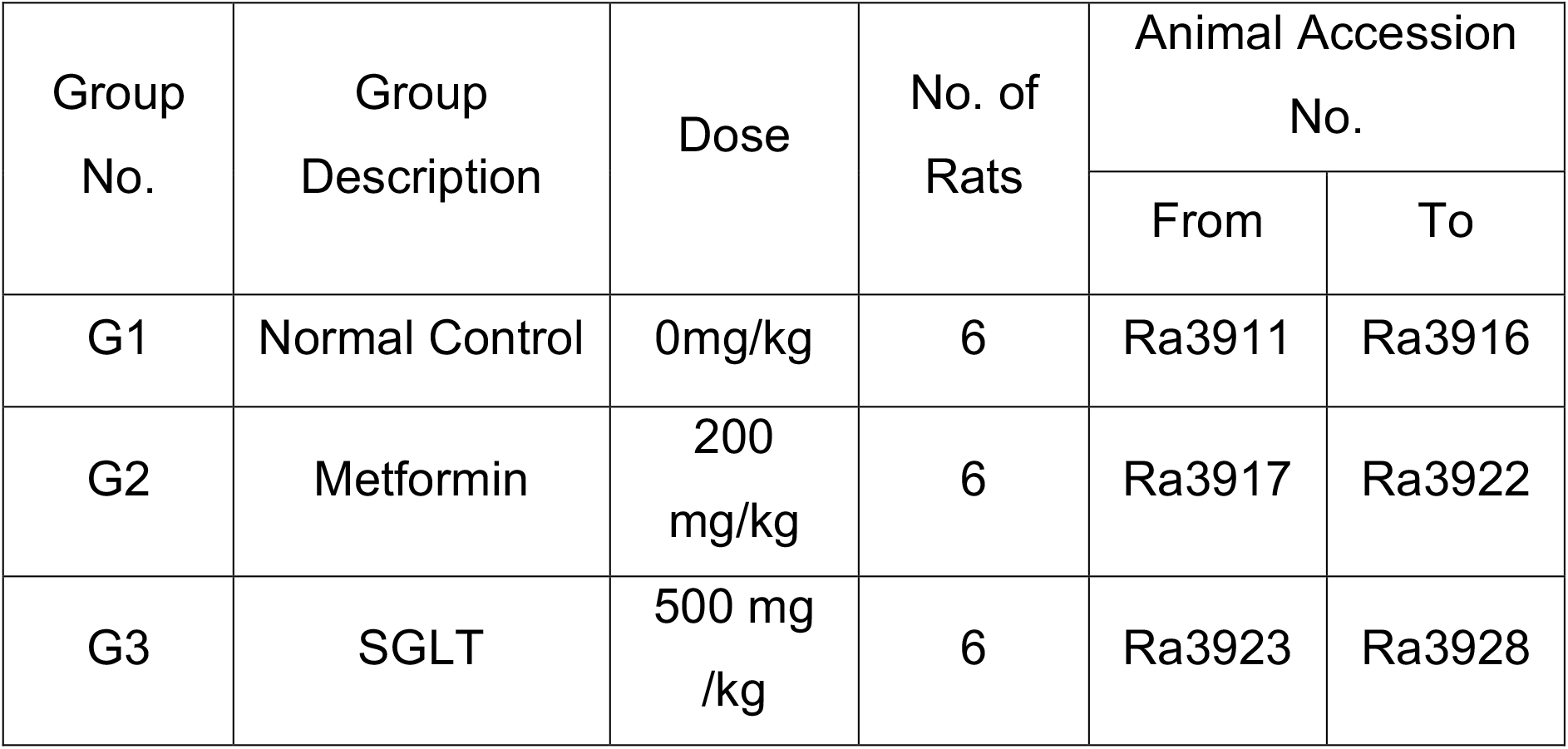

#### Test Procedure

Wistar Rats were randomly divided into 3 groups. Single administration of vehicle (Normal saline) and test item (metformin or SGLT1 IgY antibodies) were administered for respective groups. After 30 min glucose were administered at a dose of 2gm/kg and at times 0min (before glucose administration), after glucose administration 30min, 60min, 90min, 120min and 180min was measured glucose level by using Accu Chek glucometer. Oral Glucose Tolerance Test (OGTT) study were conducted using overnight fasting animals. Average fasting glucose were 100-120 mg/dl.

### OBSERVATIONS

#### Mortality, Morbidity and Clinical Signs

The animals were observed for mortality and morbidity twice daily i.e., once in the morning and once in the afternoon except during holidays and weekends whereas the animals were observed for mortality and morbidity once. Animals were observed once daily for clinical signs throughout the experimental period.

#### Body Weights

Individual body weights were recorded before test item administration on Day 1 and weekly once for all groups of rats during the treatment period.

### STATISTICAL ANALYSIS

The data was subjected to statistical analysis using specific computer programmed software and the data was analysed using One-way ANOVA followed by Dunnett’s test to compare treatment group with respective group.

All analyses and comparisons were evaluated at the 5% (p<0.05) level. Statistically significant differences (p<0.05), indicated by the tests was designated throughout the report

### STUDY COMPLIANCE

The study was performed as per the mutually agreed Study Plan and the Standard Operating Procedures (SOPs) of the Test Facility. The use of animals for this study had been approved by Liveon Biolabs Private Limited Institutional Animal Ethics Committee (IAEC). IAEC approved protocol number LBPL-IAEC-014-01/2021. Liveon Biolabs Private Limited is an AAALAC International accredited facility and registered with CPCSEA, Department of Animal Husbandry and Dairying (DAHD), Ministry of Fisheries, Animal Husbandry and Dairying (MoFAH&D), Government of India. Also, Liveon Biolabs Private Limited ensures that animal experiments are performed in accordance with the recommendation of the regulatory guidelines for laboratory animal facility published in the gazette of India, 2021.During the conduct of study none of the animals were injured, and no moribund animals were observed.

## Results

### 1. Identification of SGLT1 sequence to generate antibodies

The extracellular glucose binding domain in the SGLT1 protein has been identified and its functional role in glucose transport in the gut and kidney has been delineated. This portion of glucose binding domain can be exploited to antagonize the binding of glucose to SGLT1 and thus prevent dietary glucose absorption. We propose to design antibodies to this part of the extracellular domain as antagonists. Specifically, we will raise the IgY antibodies in an avian system.

The Basic Local Alignment Search Tool (BLAST) finds similar /conserved amino acid sequences in the same protein in various animal species. We used BLAST to compare the conserved sequences in extracellular glucose binding portion of SGLT proteins in human as well as other animal species. We identified a 19 amino acid sequence which appears to be the most conserved amino acid stretch in the extracellular glucose binding domain amongst human and animal species suggesting it may be critical in glucose uptake (Figure 2). Using these 19 amino acid containing peptide we immunized chickens and harvested IgY antibodies from the eggs.

**Figure 2.**
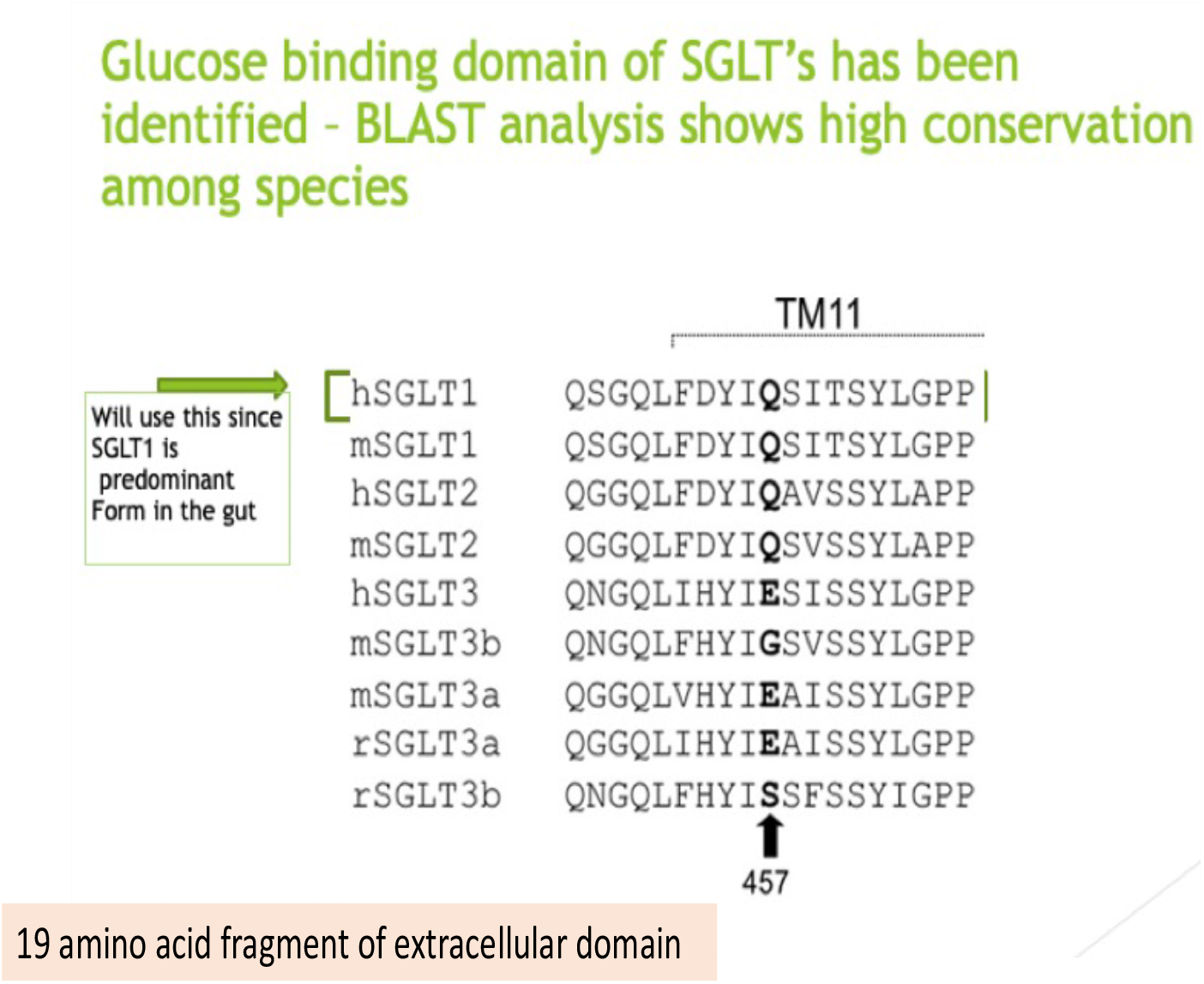

### 2. Cross Reactivity of IgY against antigen SGLT1 glucose binding domain derived peptide

Chicken IgY were generated using the peptide antigen shown above and tested for their binding specificity to the peptide using an ELISA to asses cross reactivity of the antibodies to the antigen (Figure 3). Shown below is the binding of three IgY preparation to antigen peptide coated on the plate – First batch of IgY after immunization (termed old in figure 3), second batch of IgY after immunization harvested after few days later (termed new in figure 3) and finally a non-specific IgY (anti snake venom IgY, termed snake IgY in figure 3). As shown below there was very high binding of first batch IgY against SGLT peptide antigen (absorbance OD at 540 nm in ELISA > 2) suggesting high titre of anti-SGLT IgY being elicited (Figure 3). The second batch was less cross reactive with the SGLT1 peptide antigen. In rest of the studies, the first batch of IgY were used.

**Figure 3.**
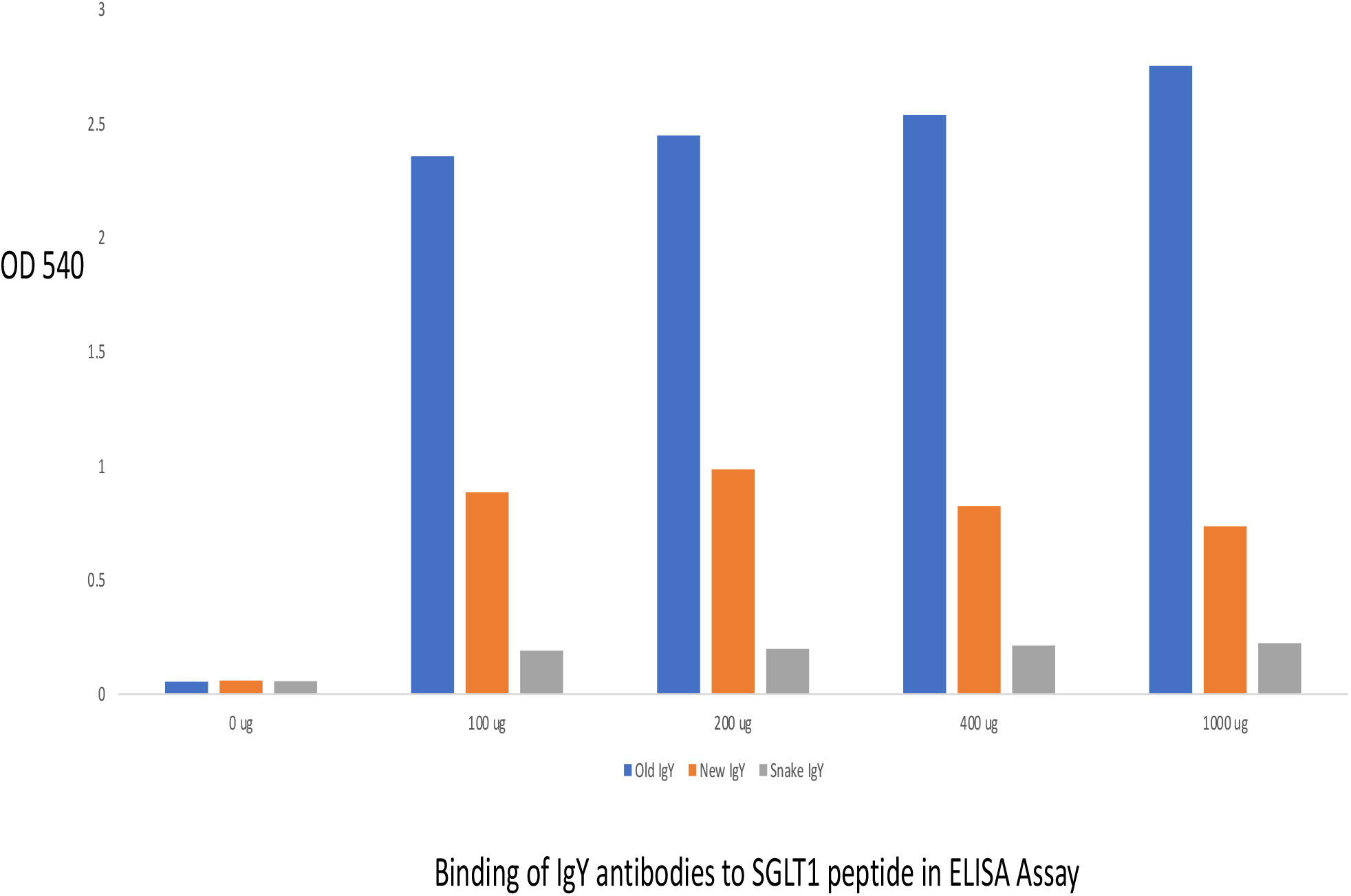

### 3. Inhibitory activity of anti SGLT1 IgY antibodies on glucose uptake in Caco2 cells

To ascertain if the anti SGLT1 antibodies are able to block glucose uptake in an intestinal human cell line, we tested the effect of various concentration of anti-SGLT1 IgY on glucose uptake in caco2 human intestinal cell line. As shown in the figure below (figure 4) there was a concentration dependent decrease in glucose uptake from 1mg/ml to 4 mg/ml with near complete inhibition at the highest concentration tested. We also test the SGLT1 peptide itself to see if it would have any antagonist activity since it was designed based on the glucose binding domain of SGLT1, and it was also found to be active at the only concentration tested.

**Figure 4.**
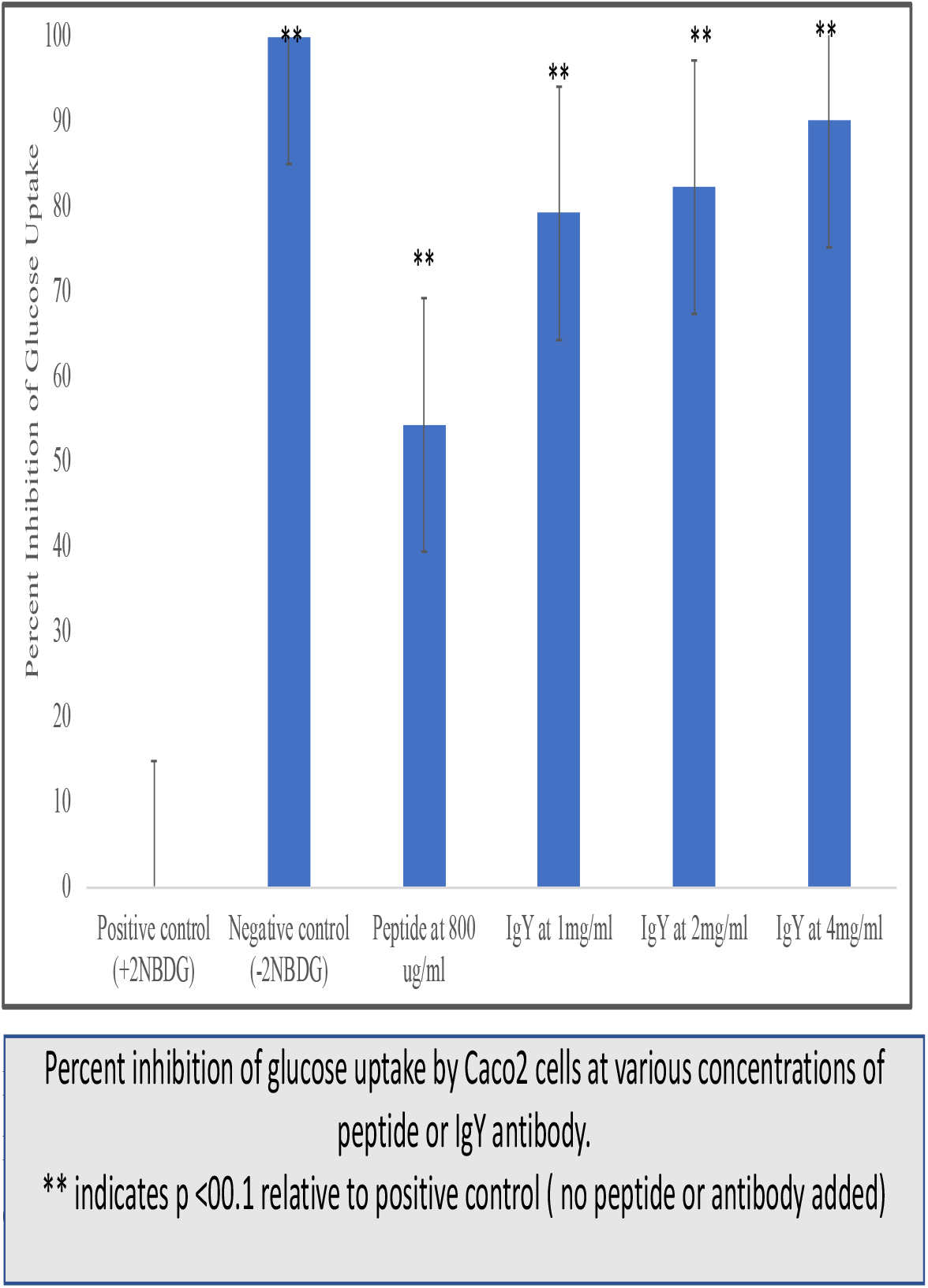

### 4. Effect of anti-SGLt1 IgY antibodies on blood glucose and glucose disposal in rat Oral Glucose Tolerance Test (OGTT)

Since the antibodies were active in blocking the uptake of glucose by Caco2 cells, i.e. functionally active, we next examined their effects in vivo on rise in blood glucose after the animals were given a bolus of glucose orally. This test is also known as OGTT and often used to detect diabetes in humans. The expectation was that if the activity is similar in vivo, to that of in vitro i.e. they block absorption/uptake of glucose by the small intestine, it should reflect in blunted rise of blood glucose levels. One group of animals received saline as pre-treatment, another received metformin given orally and a third group received oral bolus of anti-SGLT1 antibodies prior to oral glucose. After oral glucose the changes in blood glucose were monitored. As shown in Table 1, at every time point starting at 30 minutes until 180 minutes (last time point monitored) blood glucose levels in anti-SGLT1 treated animals were significantly lower and comparable to the reduction in blood glucose by metformin a drug known to block intestinal glucose absorption. The data are also shown as line graph below (Figure 5). These data confirm that the anti-SGLT1 antibodies are functionally active in vivo and demonstrate a meaningful decrease in ascending blood glucose levels after oral glucose load.

**Table 1.**
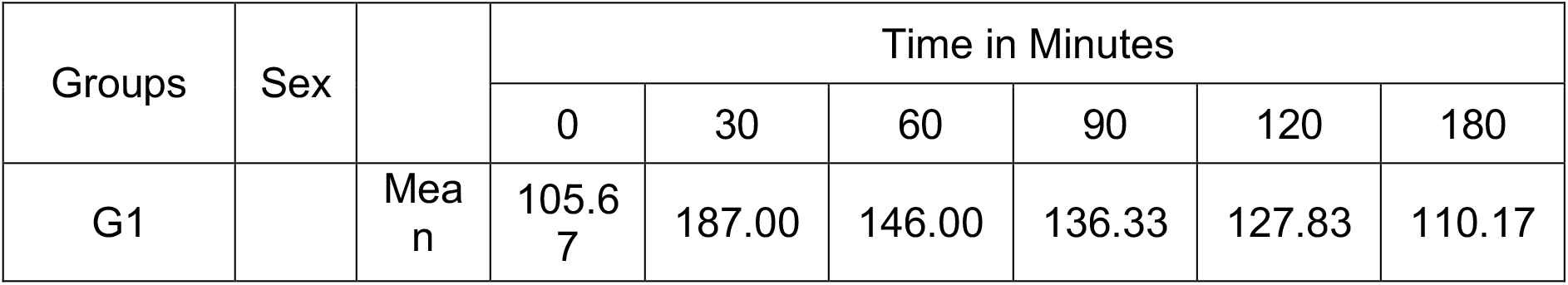
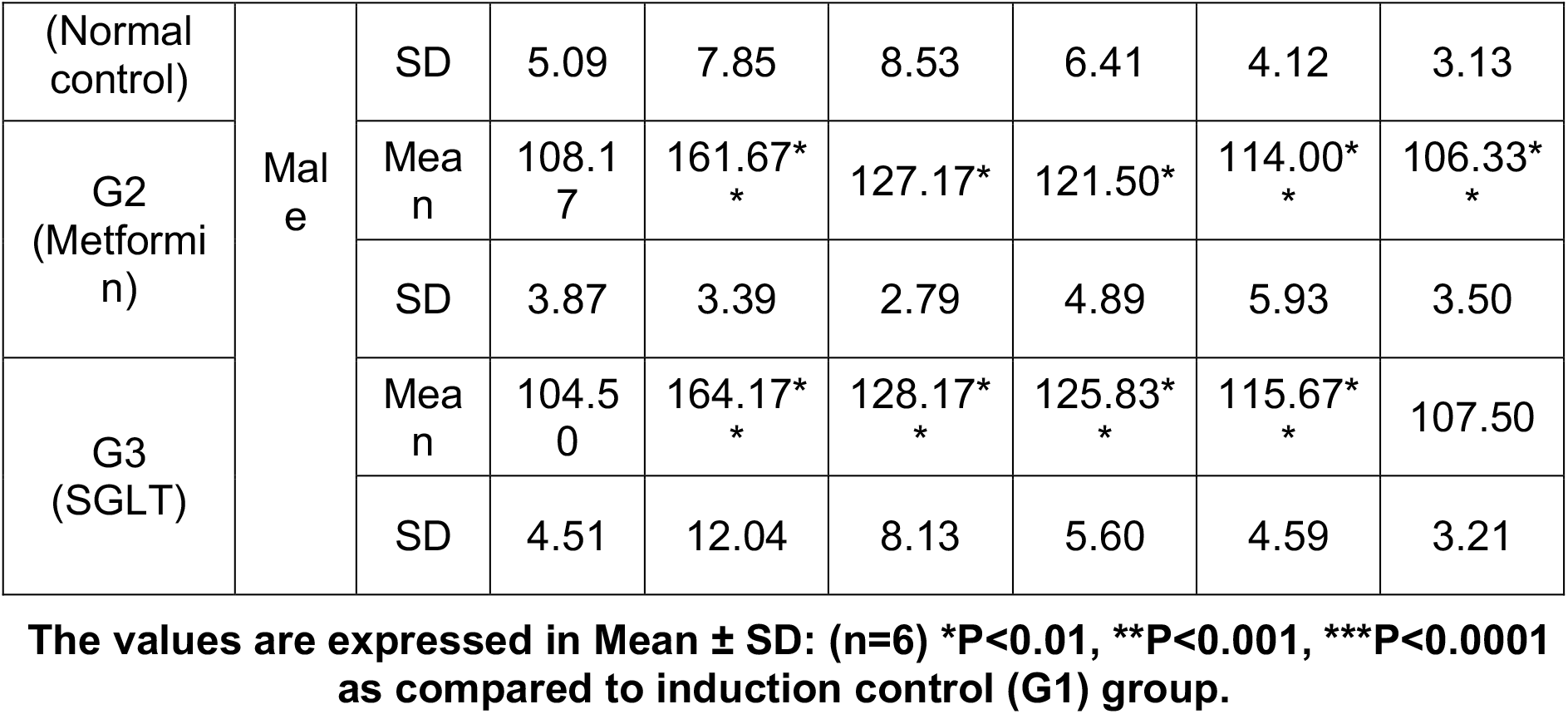
Summary of blood glucose level values (mg/dL) (Single dose OGTT study)

**Figure 5.**
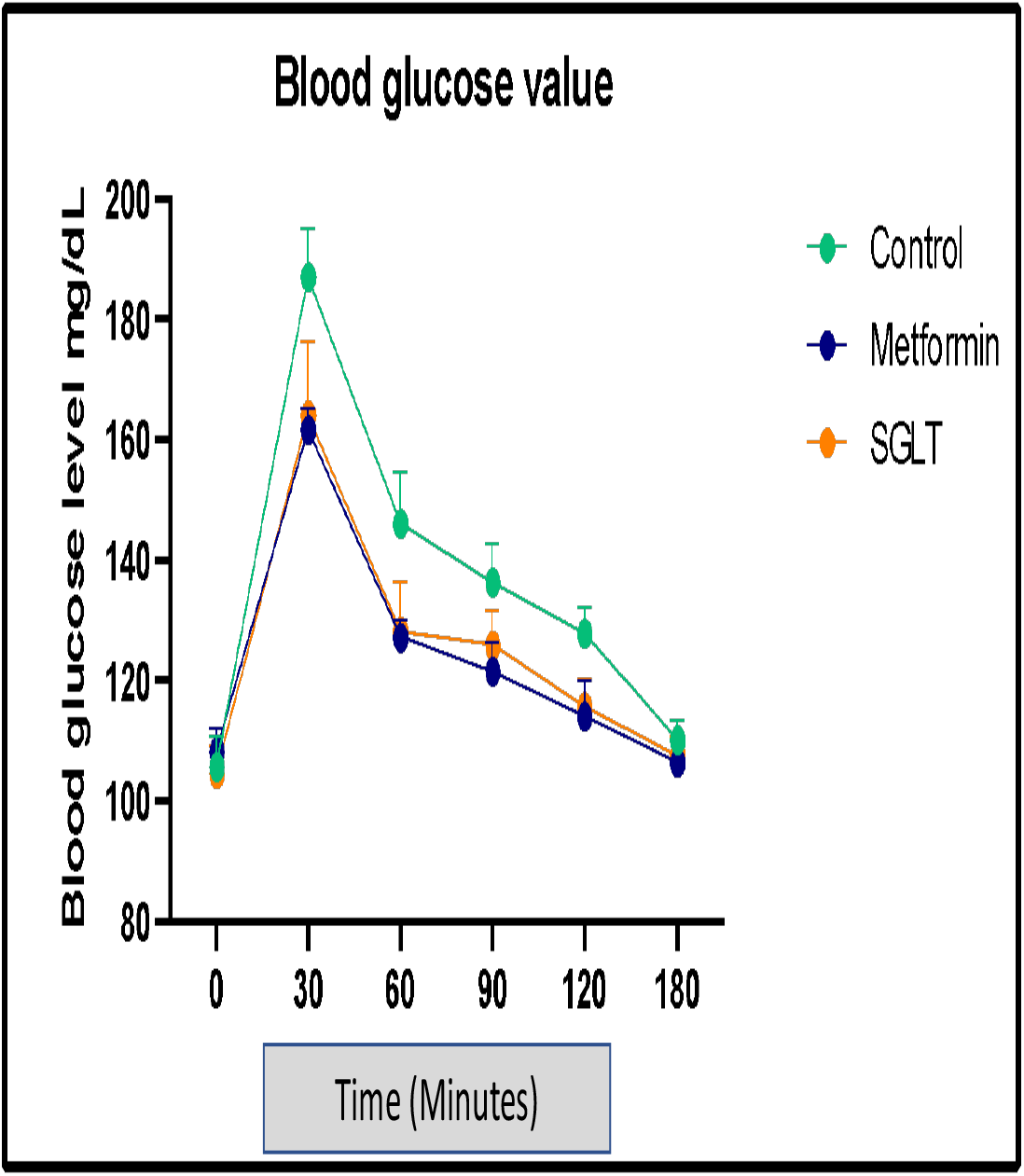

We also calculated and plotted the AUC (area under the curve) of blood glucose in each group, an indication of total glucose exposure levels after oral glucose load. A reduction in glucose exposure is indicative of lower glucose intestinal absorption and/or enhanced clearance of glucose from circulation. As shown in Table 2 below, the AUC in animals treated with metformin or anti-SGLT1 antibodies was significantly reduced suggesting an overall improved glycemic profile. The line graph of AUC data is shown in Figure 6.

**Table2.**
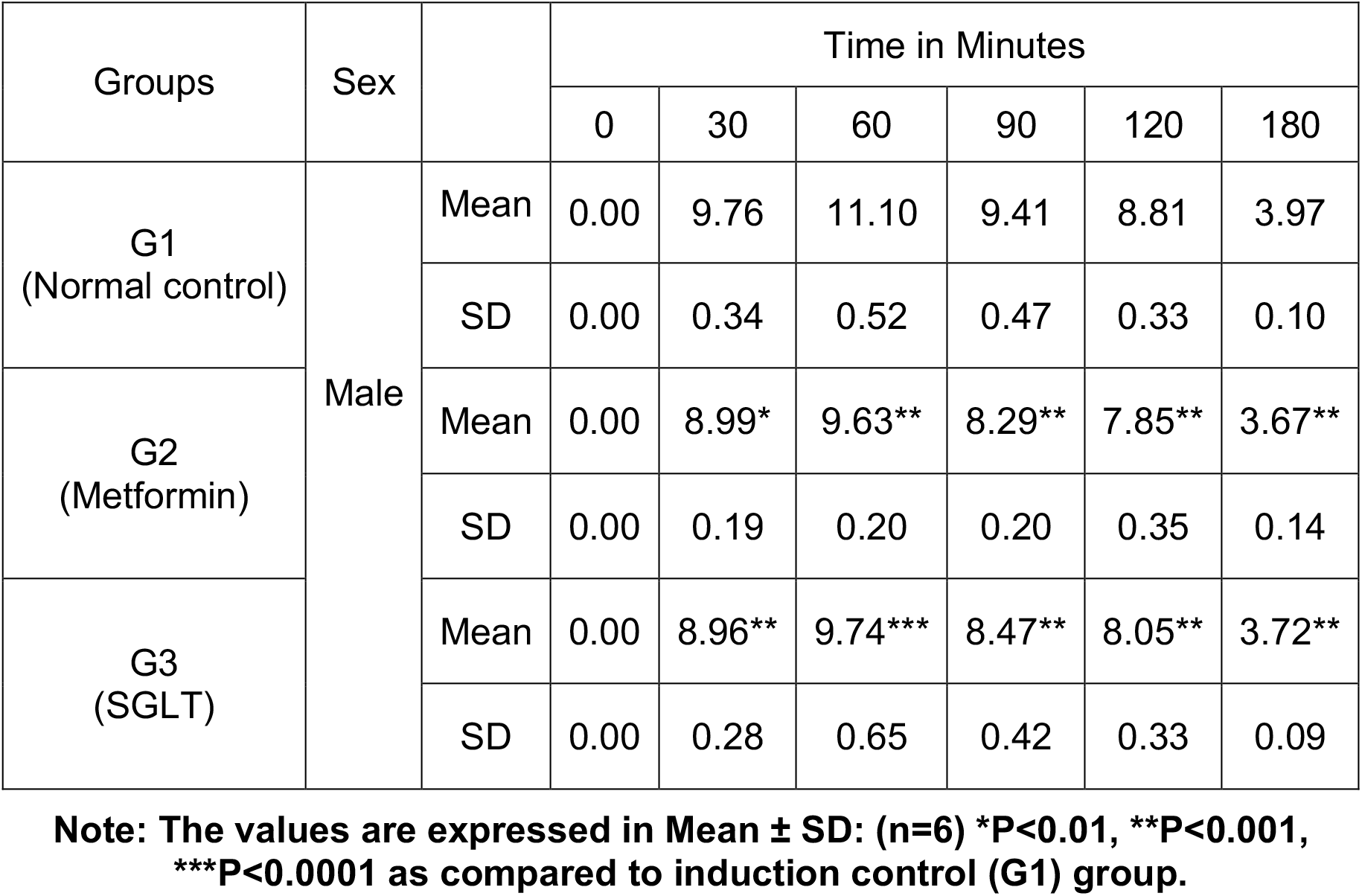
Summary of area under curve values (mg/dL) (Single dose OGTT study)

**Figure 6.**
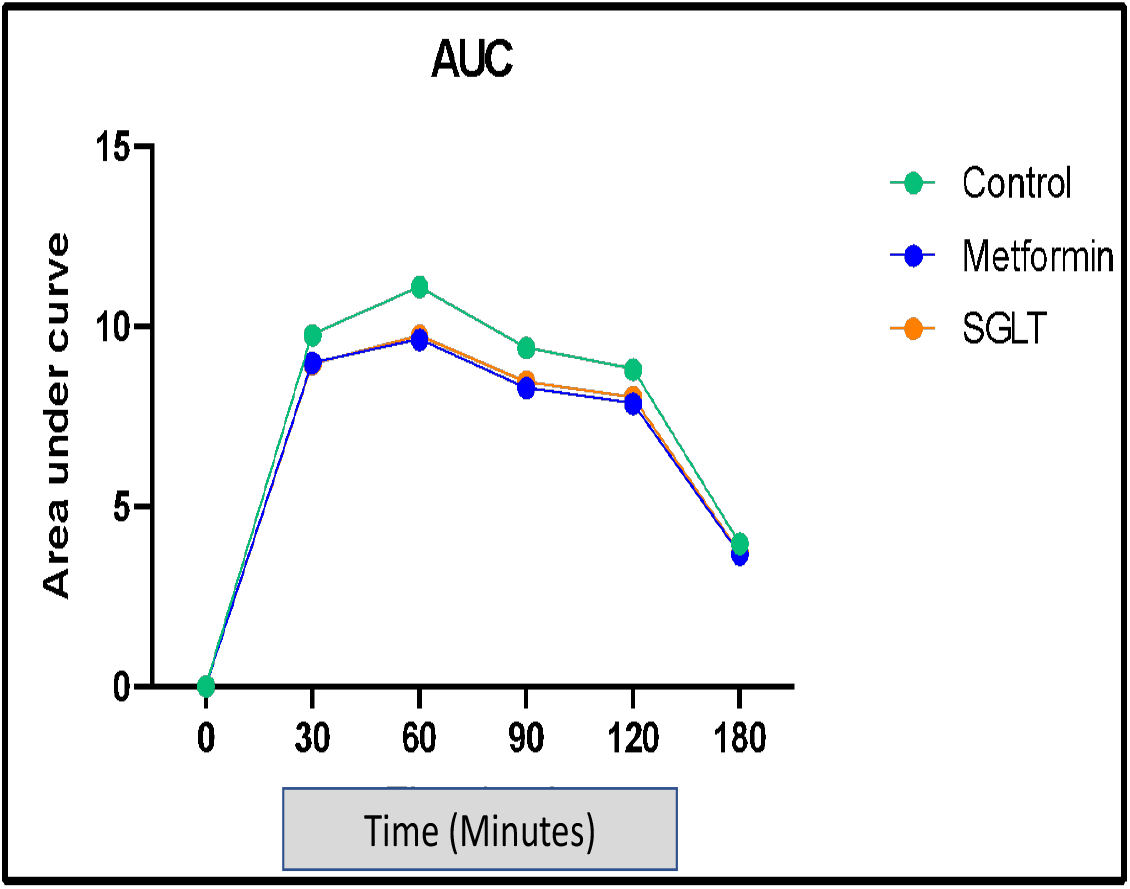

## Discussion

Our strategy was to raise IgY against the conserved 19 amino acid region of SGLT1 extracellular glucose binding domain and use the antibodies orally. We show here that oral dosing of anti-SGLT1 IgY antibodies are able to block glucose uptake and improve overall glycemic profile in vivo in OGTT studies. Since egg-derived chicken IgY are considered GRAS (generally regarded as safe) and safe for human consumption we could use these as oral over the counter product or a prescription product for treatment of hyperglycaemia.

SGLT-1 is selectively expressed in intestine and other limited organ set, thus making it an attractive target for manipulation of glucose metabolism. Dietary glucose absorption begins in the buccal cavity itself and extends to the small intestine – therefore a therapy that can block glucose absorption starting in the mouth such as ant-SGLt1 antibodies and the intestine would be beneficial.

Currently patients with hyperglycemia/diabetes of any kind are put on a regimen of diet, exercise and other medications. Yet reaching target glucose levels is still not possible for many due to intolerance to some of the drugs. The additional advantage of our approach is that the therapy is GRAS and can easily be added as an adjunct to existing regimen. This could provide a new option for the physicians in treatment of this disease including gestational diabetes where drug treatment options are very limited.

